## Supplementary Information for "Cryo-EM structures of human magnesium channel MRS2 reveal gating and regulatory mechanisms"

Supplementary Fig. 1 Protein purification, biochemical characterization, and negative-staining EM analysis of human MRS2.
Supplementary Fig. 2 Single-particle cryo-EM data processing of MRS2-$Mg^{2+}$.
Supplementary Fig. 3 Density fit of the MRS2-$Mg^{2+}$ model.
Supplementary Fig. 4 Density fit of $Mg^{2+}$-binding sites, gating residues, and the R116-E291 pair in the MRS2-$Mg^{2+}$ model.
Supplementary Fig. 5 Structural comparison between *Hs*MRS2 and CorA superfamily members.
Supplementary Fig. 6 Sequence alignment of MRS2 and CorA.
Supplementary Fig. 7 Structural comparison between MRS2-$Mg^{2+}$ and MRS2-EDTA, and densities near the soluble $Mg^{2+}$-binding sites 1 and 2.
Supplementary Fig. 8 Density near the R332-ring in the MRS2-$Mg^{2+}$ and MRS2-EDTA structures.
Supplementary Fig. 9 Overview of proposed models of Mg2+-translocation mechanisms in MRS2 and CorA family members

Supplementary Table 1 Cryo-EM data collection parameters and analysis.
Supplementary Table 2 Cryo-EM map and model analysis.

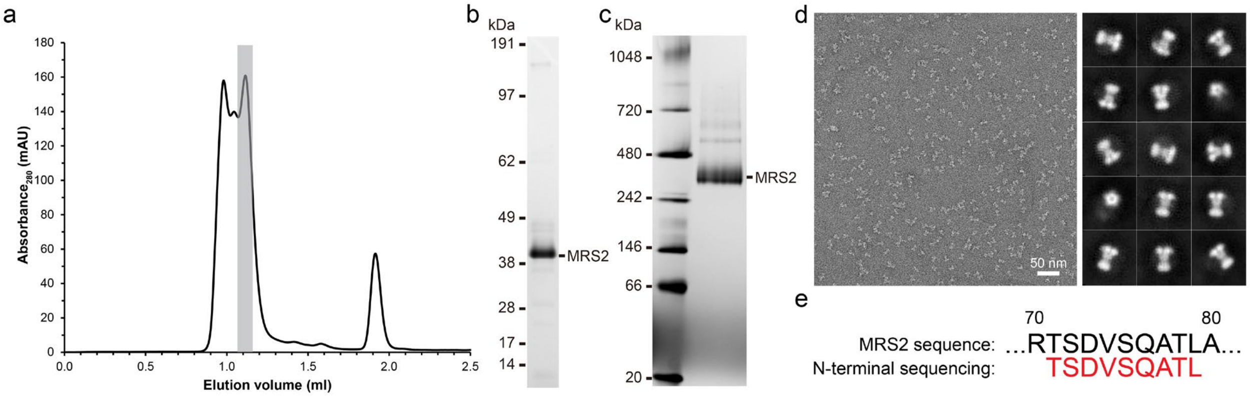

**Supplementary Fig. 1 Protein purification, biochemical characterization, and negative-staining EM analysis of human MRS2**. **a**, Size-exclusion chromatography elution profile of MRS2 obtained after M2 agarose affinity chromatography. Fractions highlighted were analyzed by Coomassie-stained SDS-PAGE **(b)**, BN-PAGE **(c),** and negative-staining EM **(d)**. **b-c**, SDS-PAGE gel **(b)** and BN-PAGE gel **(c)** of purified MRS2. **d**, Negative-staining EM imaging of purified MRS2 in the presence of $Mg^{2+}$. Representative micrograph (left) and the selected 2D class averages (right) with a cropped box size of 90 px (~274 Å) are shown. **e**, N-terminal sequencing detecting the first 9 amino acids of the purified human MRS2 indicated in red.

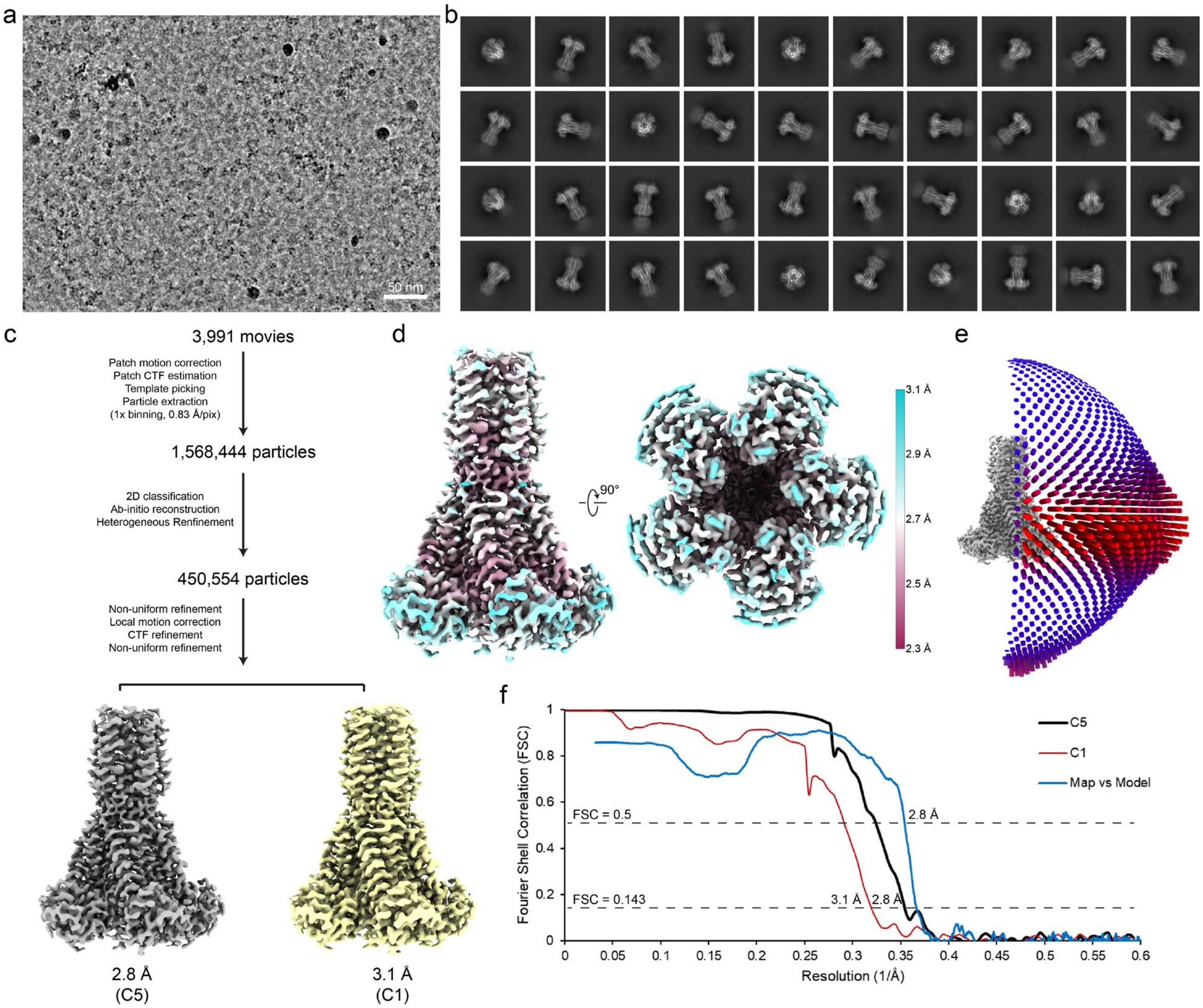

**Supplementary Fig. 2 Single-particle cryo-EM data processing of MRS2-$Mg^{2+}$. a-b**, Representative cryo-EM micrograph **(a)** and selected 2D class averages **(b)** of MRS2 with a box size of 320 px (~266 Å). **c**, Workflow of cryo-EM image processing of MRS2. The resolution was reported according to the Fourier shell correlation (FSC) = 0.143 criteria. Refinements with C5 and C1 symmetry were performed and local resolution filtered maps are shown. The map refined with C5 symmetry was used for further structural analysis throughout the current study because of higher resolution, while the C1 map was used to confirm the ion densities along the symmetry axis. **d**, Local resolution evaluation of the MRS2 density map at an average 2.8 Å resolution. **e-f**, Evaluation of the cryo-EM reconstruction of the MRS2 maps with Euler angle distribution plot **(e)** and FSC curves **(f)**.

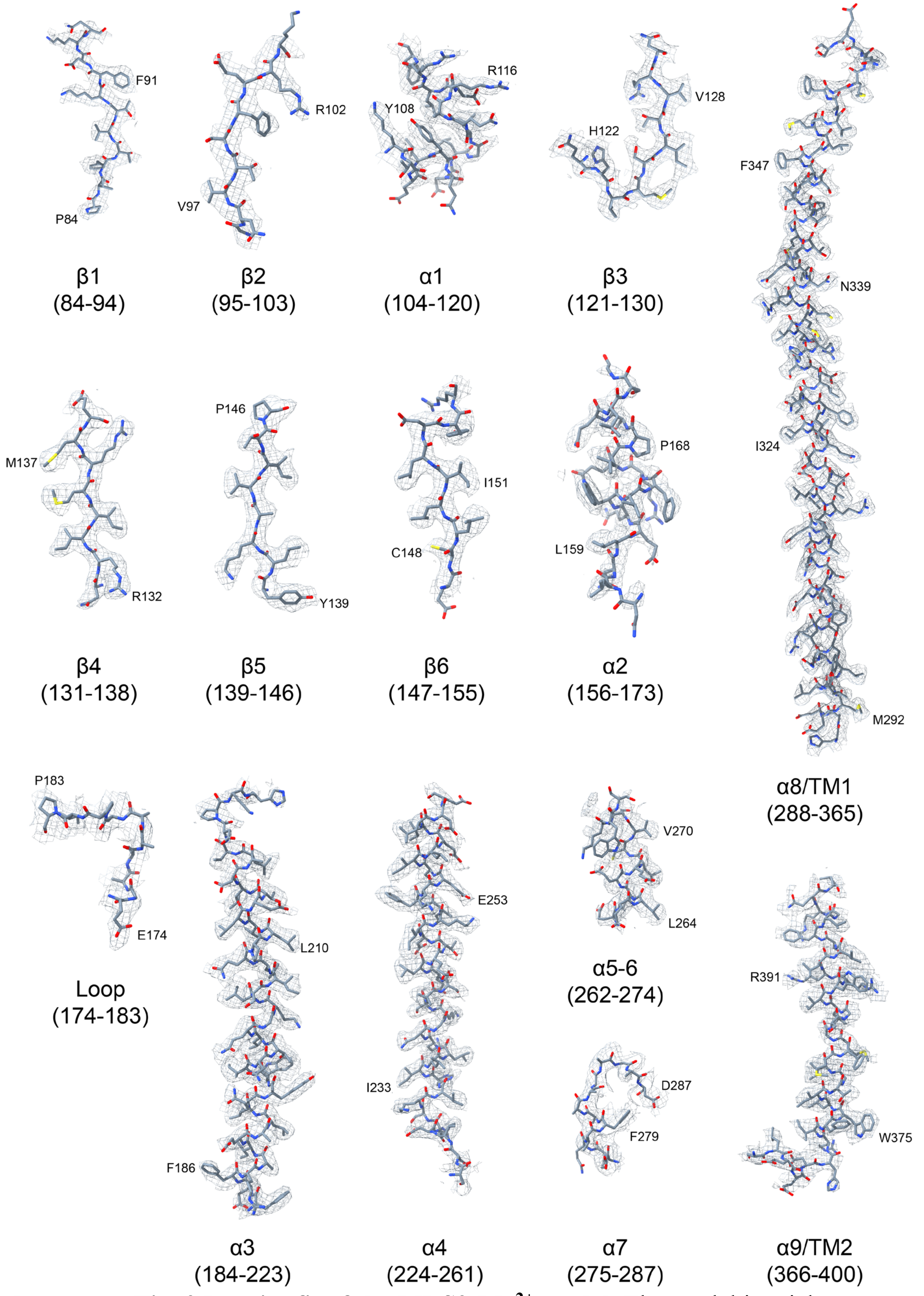

**Supplementary Fig. 3 Density fit of the MRS2-$Mg^{2+}$ model.** The model in stick representation is superimposed on the cryo-EM densities in mesh.

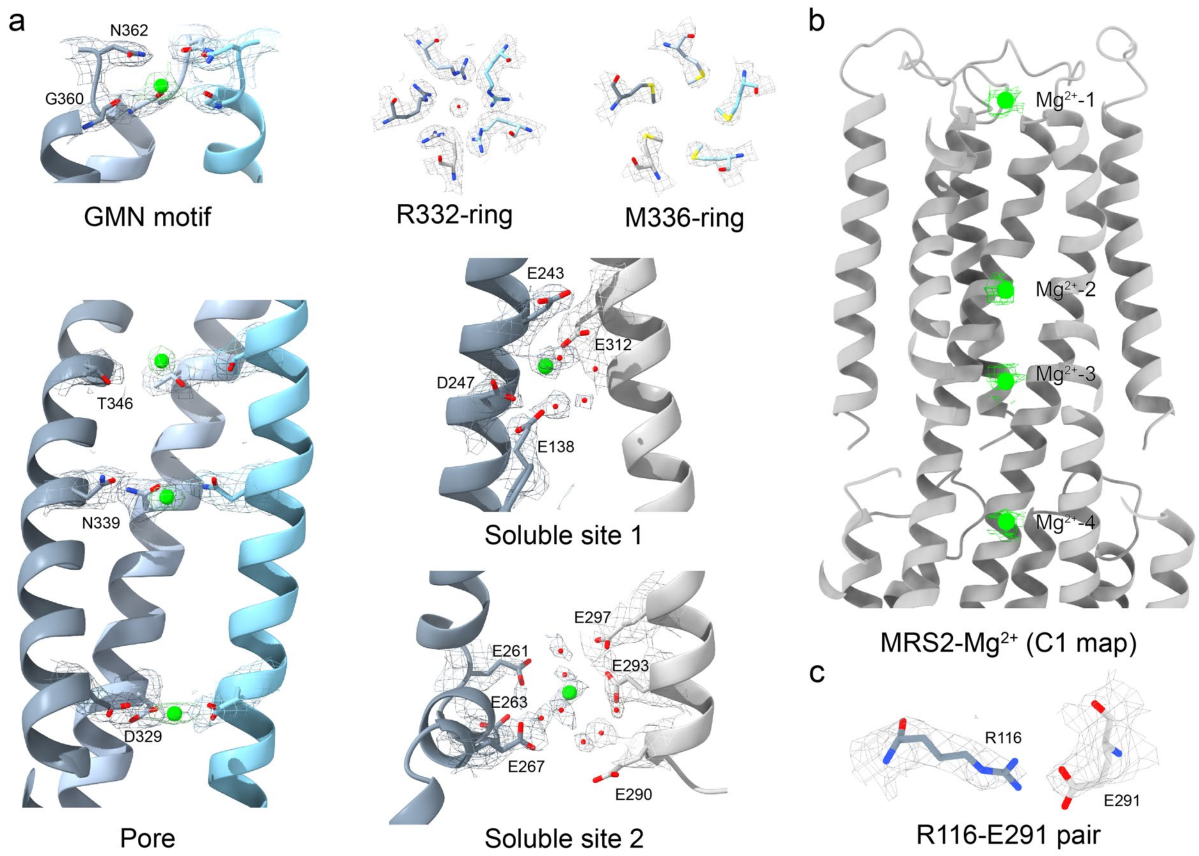

**Supplementary Fig. 4 Density fit of $Mg^{2+}$-binding sites, gating residues, and the R116-E291 pair in the MRS2-$Mg^{2+}$ model**. **a**, Densities of $Mg^{2+}$ ions, R332- and M336-ring, coordinating water molecules and residues are shown. **b**, Densities of $Mg^{2+}$ along the pore in the C1 map of the MRS2-$Mg^{2+}$ structure. **c**, Densities of the R116-E291 pair.

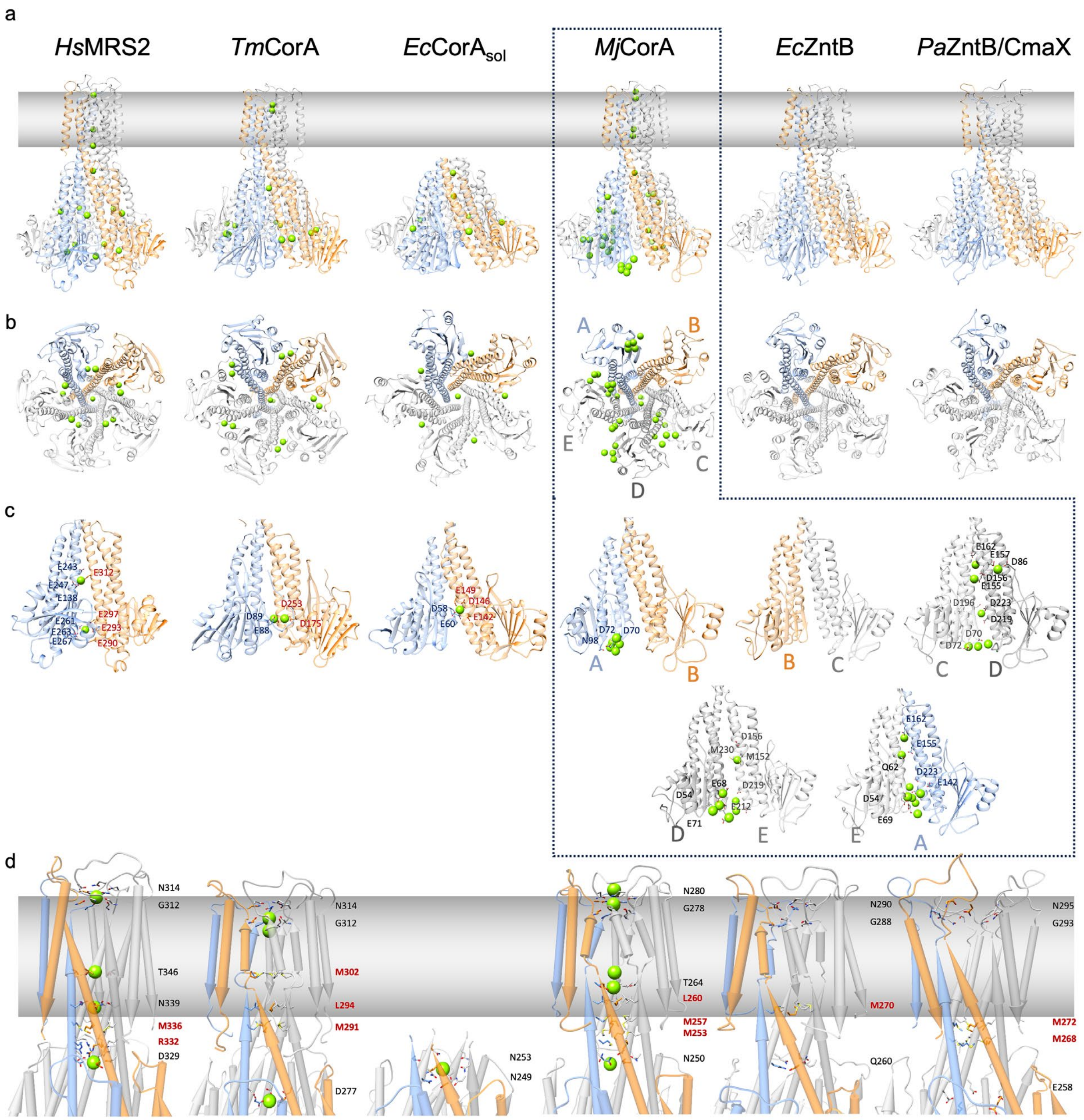

**Supplementary Fig. 5 Structural comparison between *Hs*MRS2 and CorA superfamily members. a-b**, Side **(a)** and bottom **(b)** views of *Hs*MRS2 (MRS2-$Mg^{2+}$, current study), *Tm*CorA (PDB: 3FCF), *Ec*CorA$_{sol}$ (PDB: 5N77), *Mj*CorA (PDB: 4EV6), *Ec*ZntB (PDB: 5N9Y) and *Pa*ZntB/CmaX (PDB: 7HN9) are shown with two of the five subunits highlighted in blue and orange. $Mg^{2+}$ ions are shown as green spheres. **c,** Two neighboring soluble domain subunits of these structures are shown highlighting residues near assigned $Mg^{2+}$ ions. Owing to the asymmetric arrangement of *Mj*CorA, side views of each pair are shown. **d**, Comparison between the pore regions of the corresponding structures shown in **(a)** using UCSF Chimera's Pipes and Planks representation. Residues coordinating the pore $Mg^{2+}$ are shown and labelled. Gating residues are shown and labelled in red.

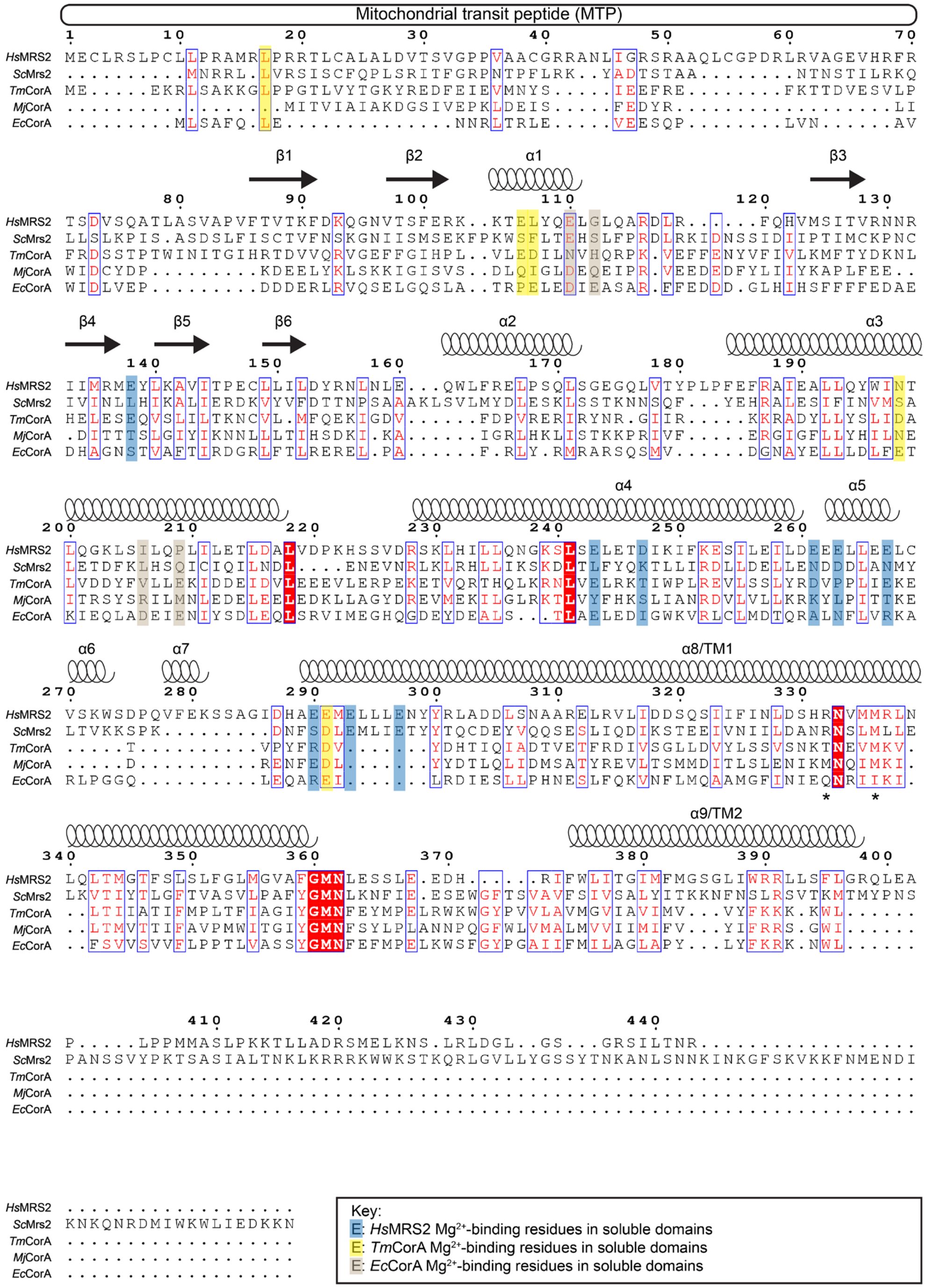

**Supplementary Fig. 6 Sequence alignment of MRS2 and CorA.** Sequences of MRS2 from *Homo sapiens*, *Saccharomyces cerevisiae*, and CorA from *Thermotoga maritima*, *Methanocaldococcus jannaschii*, and *Escherichia coli* were subjected to PRALINE multiple sequence alignment. $Mg^{2+}$-binding residues in the soluble domain of human MRS2, *Tm*CorA and *Ec*CorA are mapped and highlighted in blue, yellow, and tan, respectively. The gating residues of *Hs*MRS2 are denoted by asterisks. The mitochondrial transit peptide (MTP) and the secondary structures of human MRS2 are depicted above the sequences. Sequence identifies of *Hs*MRS2 to *Sc*MRS2, *Tm*CorA, *Mj*CorA, and *Ec*CorA are 20%, 14%, 16%, and 17%, respectively.

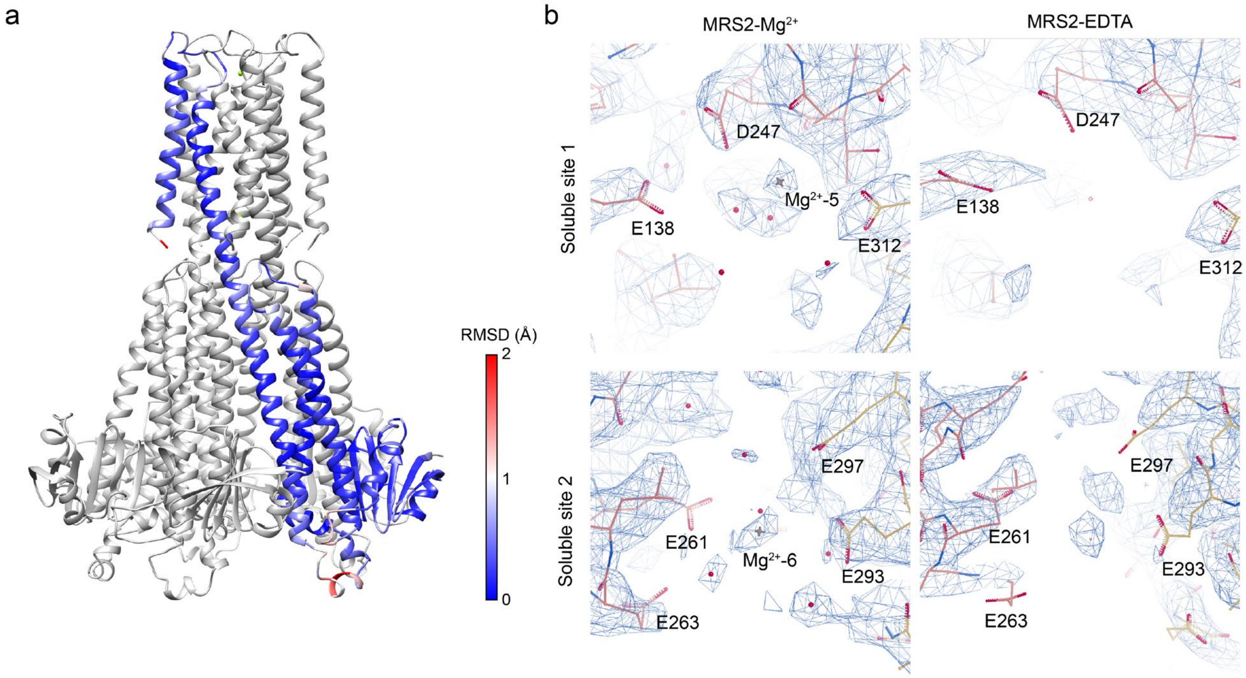

**Supplementary Fig. 7 Structural comparison between MRS2-$Mg^{2+}$ and MRS2-EDTA, and densities near the soluble $Mg^{2+}$-binding sites 1 and 2. a**, The MRS2-EDTA model with one subunit colored according to RMSD between the MRS2-$Mg^{2+}$ and MRS2-EDTA structures. **b**, Densities near the soluble $Mg^{2+}$-binding sites 1 and 2 of MRS2-$Mg^{2+}$ (left) and MRS2-EDTA (right).

MRS2-Mg$^{2+}$

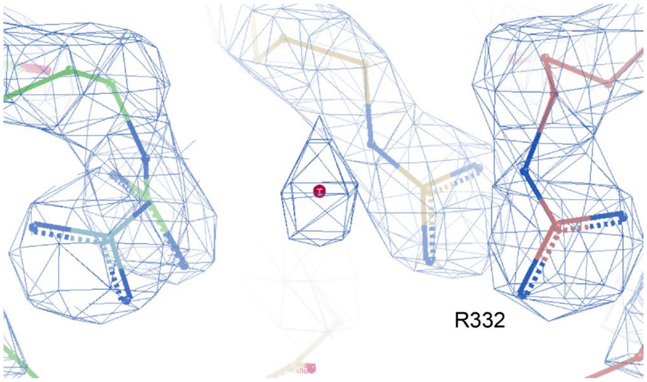

MRS2-EDTA

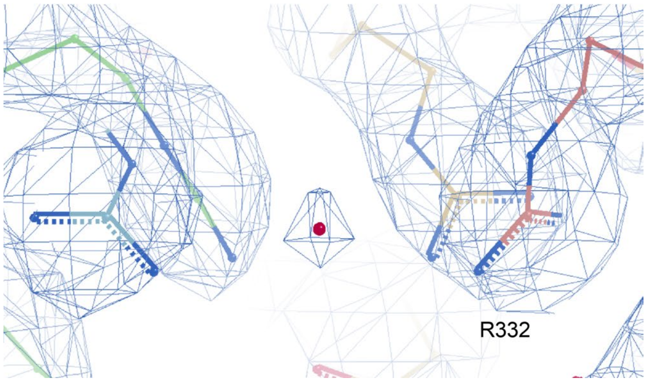

**Supplementary Fig. 8 Density near the R332-ring in the MRS2-Mg$^{2+}$ and MRS2-EDTA structures.** Fitted structures in the MRS2-Mg$^{2+}$ (upper) and MRS2-EDTA (lower) cryo-EM density maps in mesh. The extra density is also observed in the non-symmetrized MRS2-Mg$^{2+}$ C1 map.

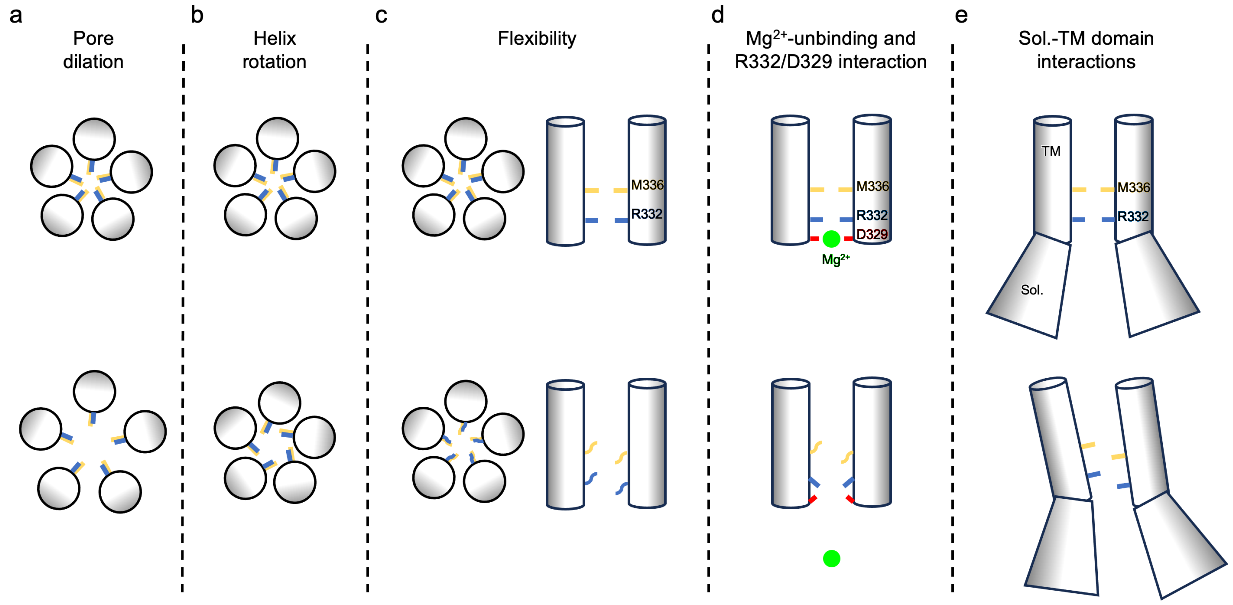

**Supplementary Fig. 9 Overview of proposed models of $Mg^{2+}$-translocation mechanisms in MRS2 and CorA family members.** Based on all known structures of CorA family members to date, the pore has been observed to be too narrow to allow hydrated magnesium ions to pass through. An early model proposed that the pore needs to dilate or expand **(a)** to allow for translocation of $Mg^{2+}$ which has been observed in other channels that have been structurally resolved in a closed and more expanded open conformation. Another proposed model relies on a conformational change in form of a helix rotation **(b)** which would alter the pore facing residues. A third model is based on flexibility **(c)** which would ultimately allow for ion translocation. Here, we propose that the $Mg^{2+}$ bound to the D329-ring plays a role in regulating the R332 gate **(d)**. Under high matrix $Mg^{2+}$ concentration, the presence of $Mg^{2+}$ around D329-ring decouples the interaction between D329 and the R332 gate. When the matrix $Mg^{2+}$ concentration is low, unbinding of this $Mg^{2+}$ occurs, which allows D329 to interact with R332, thereby lowering the energy barrier for $Mg^{2+}$ permeation. In addition, a connection between gating residue R332 and the soluble domain of an adjacent subunit via hydrogen bonds is observed. Change in conformation of the soluble domains may couple these with TM movement leading to gate opening **(e)**.

**Supplementary Table 1 Cryo-EM data collection parameters and analysis.**

| | **MRS2-$Mg^{2+}$** | **MRS2-EDTA** |
|---|---|---|
| Date of collection | 2022/05/9-11 | 2022/5/12-15 |
| Protein concentration | 0.5 mg/mL | 0.5 mg/mL |
| Sample volume | 3 µl | 3 µl |
| Grid type | QF R 1.2/1.3 400 Cu mesh | QF R 1.2/1.3 400 Cu mesh |
| Plunge freezer | Leica EM GP2 | Leica EM GP2 |
| Blotting time (s) | 6 | 7 |
| Temperature (°C) | 4 | 4 |
| Humidity (set) | 95% | 95% |
| Microscope | Titan Krios G1 | Titan Krios G1 |
| Voltage (kV) | 300 | 300 |
| Camera | K3 (CDS mode) | K3 (CDS mode) |
| Energy filter (slit) | Yes (20 eV) | Yes (20 eV) |
| Cs corrector | No | No |
| Objective aperture | 70 µm | 70 µm |
| Magnification | 105,000x | 105,000x |
| Physical pixel size (Å/px) | 0.83 | 0.83 |
| Super-resolution pixel size (Å/px) | 0.415 | 0.415 |
| Electron exposure ($e^-/Å^2$) | 50 | 50 |
| Number of movie frames | 50 | 50 |
| Dose rate ($e^-$/px/s) | 10 | 10 |
| Defocus (µm) | -0.7 to -2 | -0.7 to -2 |
| Number of total micrographs | 3,991 | 9,659 |
| Number of selected micrographs | 3,286 | 9,238 |
| Number of particles picked | 1,568,444 | 4,802,706 |

**Supplementary Table 2 Cryo-EM map and model analysis.**

| | MRS2-$Mg^{2+}$ (C5) EMD-41624 PDB ID 8TUL | MRS2-$Mg^{2+}$ (C1) EMD-41629 | MRS2-EDTA (C5) EMD-41628 PDB ID 8TUP | MRS2-EDTA (C1) EMD-41630 |
|---|---|---|---|---|
| **Data processing** | | | | |
| Final number of particles | 450,544 | 450,544 | 1,744,117 | 1,744,117 |
| Final pixel size used for final maps (Å/px) | 0.83 | 0.83 | 0.83 | 0.83 |
| Symmetry imposed | C5 | C1 | C5 | C1 |
| Resolution of map (Å) | 2.8 | 3.1 | 3.3 | 3.6 |
| B-factor for map ($Å^2$) | 141 | 119 | 201 | 181 |
| **Model composition** | | | | |
| Non-hydrogen atoms | 13,070 | | 12,869 | |
| Water | 281 | | 66 | |
| $Mg^{2+}$ ions (Ligand) | 14 | | 3 | |
| **Refinement and validation** | | | | |
| Model resolution (Å) | 2.8 | | 3.3 | |
| FSC threshold | 0.5 | | 0.5 | |
| Bond lengths RMSD (Å) | 0.003 | | 0.002 | |
| Bond angle RMSD (°) | 0.472 | | 0.405 | |
| MolProbity score | 1.58 | | 1.28 | |
| Clashscore | 5.94 | | 4.52 | |
| Rotamer outliers (%) | 2.14 | | 0.00 | |
| Ramachandran outliers (%) | 0.00 | | 0.00 | |
| Ramachandran favored (%) | 98.10 | | 98.10 | |
| Ramachandran allowed (%) | 1.27 | | 2.22 | |
